# Active Transport by Cytoplasmic Dynein Maintains the Localization of MAP2 in Developing Neurons

**DOI:** 10.1101/2023.04.26.538370

**Authors:** Yoji Yonemura, Yuri Sakai, Rinaho Nakata, Ayaka Hagita-Tatsumoto, Tomohiro Miyasaka, Hiroaki Misonou

## Abstract

MAP2 has been widely used as a marker of neuronal dendrites because of its extensive restriction in the somatodendritic region of neurons. Despite that, how the precise localization of such a soluble protein is established and maintained against thermal forces and diffusion has been elusive and long remained a mystery in neuroscience. In this study, we aimed to uncover the mechanism behind how MAP2 is retained in the somatodendritic region. Using GFP-tagged MAP2 expressed in cultured hippocampal neurons, we discovered a crucial protein region responsible for the localization of MAP2, the serine/proline-rich (S/P) region. Our pulse-chase live-cell imaging revealed the slow but steady migration of MAP2 toward distal dendrites, which was not observed in a MAP2 mutant lacking the S/P region, indicating that S/P-dependent transport is vital for the proper localization of MAP2. Furthermore, our experiments using an inhibitor of cytoplasmic Dynein, ciliobrevin D, as well as Dynein knockdown, showed that cytoplasmic Dynein is involved in the transport of MAP2 in dendrites. We also found that Dynein complex binds to MAP2 through the S/P region in heterologous cells. Using mathematical modeling based on experimental data, we confirmed that an intermittent active transport mechanism is essential. Thus, we propose that the cytoplasmic Dynein recruits and transports free MAP2 toward distal dendrites, thereby maintaining the precise dendritic localization of MAP2 in neurons. Our findings shed light on the previously unknown mechanism behind MAP2 localization and provide a new direction for soluble protein trafficking research in the field of cell biology of neurons.

## INTRODUCTION

MAP2 is one of microtubule (MT) associated proteins (MAPs) and has been known to localize predominantly to the cell body and dendrites (Bernhardt and Matus, 1984; De Camilli et al., 1984). Among MAPs, MAP2 and Tau are probably the best-recognized and -utilized MAPs in neurobiology, owing to their usefulness as markers of neuronal dendrites and the axon, respectively. Primary function of MAP2, like Tau, is thought to bind to MT to stabilize them (Dehmelt and Halpain, 2005). Alternative splicing of MAP2 gives rise to 4 isoforms, MAP2A and B (high molecular weight, HMW), and C and D (low molecular weight, LMW). Expression of these isoforms are differently regulated during neuronal development. MAP2B is expressed throughout development, and the expression level of MAP2A increases during development. In contrast, expression level of MAP2C and D decreases during development, although LMW MAP2 still exist in adulthood (Farah and Leclerc, 2008). A physiological role of somatodendritic localization of MAP2, particularly near the hillock of the axon, has recently been proposed as a facilitator of the axonal cargo sorting before entering the axon initial segment (Gumy et al., 2017). Despite the utility and significance of the somatodendritic localization of MAP2, still very little is known about how the specific localization is established and maintained.

There have been attempts to understand how MAP2 localizes to soma and dendrites and not to the axon. We reckon following three major hypotheses proposed over the years; i) preferential binding of MAP2 to dendritic MT (Kanai and Hirokawa, 1995), ii) localization of MAP2 mRNA in dendrites for local translation (Garner et al., 1988), and iii) rapid turnover of MAP2 in the axon (Hirokawa et al., 1996). Furthermore, Kanai et al. (Kanai and Hirokawa, 1995) reported that projection domain (PD) of MAP2, which exist only in MAP2A and B, prevent MAP2 from entering the axon, resulting in the somatodendritic localization of HMW MAP2, although vast majority of LMW MAP2 still localizes to the soma and dendrites. Despite these efforts, there has not been a convincing model, which can account for the highly restricted localization of MAP2.

In this study, we sought to identify molecular mechanisms of somatodendritic localization of MAP2. We investigated factors involved in regulating the localization of MAP2 and found that the serine/proline rich region (S/P region) of MAP2 is critical for its specific localization to the soma and dendrites. Live-cell pulse-chase imaging using a photoconvertible fluorescent protein, Dendra2, revealed that MAP2 is transported toward the tip of a dendrite, and that this process is dependent on the S/P region. Furthermore, inhibiting cytoplasmic Dynein resulted in the disappearance of the dendritic transport phenotype and axonal leakage of endogenous MAP2, suggesting that Dynein transports MAP2 through the S/P region and localizes MAP2 to the somatodendritic compartment. We then built a mathematical model to test whether this mechanism is sufficient to establish and maintain the somatodendritic localization of MAP2. Simulation using Virtual Cell (VCell) (Cowan et al., 2012) demonstrated that the active transport mechanism is essential for the localization. Based on these results we propose a new model for the somatodendritic-specific localization of MAP2, which has not been unequivocally shown for decades.

## RESULTS

### Important protein domain of MAP2 for the somatodendritic localization

To understand how the exclusive dendritic localization of MAP2 is established and maintained, we analyzed human MAP2C tagged with AcGFP (GFP hereafter) transfected into cultured neurons at 7 days *in vitro* (DIV) and analyzed at 9 DIV. We found that GFP-MAP2C exhibited somatodendritic localization (Fig. 1A) despite its 3.5 ±0.5-fold overexpression (Fig. 1B). It should also be noted that overexpression of GFP-MAP2C did not result in any discernable difference in endogenous MAP2 expression nor localization (Fig. 1C). The somatodendritic localization of GFP-MAP2C indicates that protein domains, other than the PD found only in the HMW MAP2s, contains a signal for the somatodendritic retention. To identify the critical protein domains, we generated MAP2C-Tau chimera, in which the C-terminal half containing the microtubule-binding domain (MTBD) was swapped with that of Tau (Fig. 1D). Interestingly, this chimera localized like the wild-type MAP2C, as evidenced by the rapid decrease of fluorescence signals from the soma into the axon (Fig. 1E). These results suggest that the N-terminal half of MAP2C contains the critical domains for localization.

**Figure 1.**
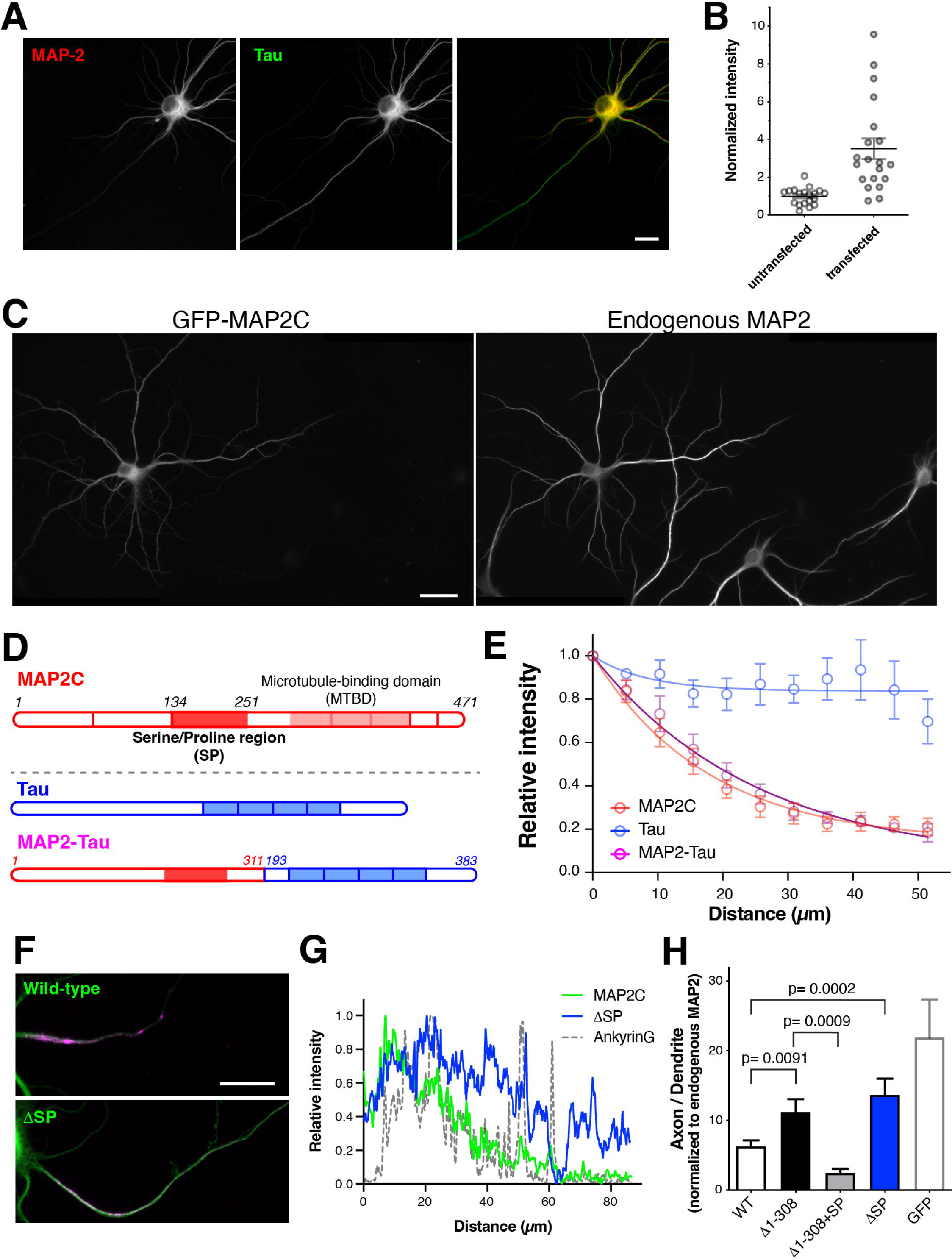
Determinant of the somatodendritic localization of MAP2C. **A,** Somatodendritic localization of GFP-tagged human MAP2C expressed in cultured neurons with mCherry-tagged human Tau. Scale bar, 20 µm. **B,** Estimation of expression levels of GFP-MAP2C using immunofluorescence staining with anti- MAP2 antibody, which detects both endogenous and exogenous MAP2 isoforms. Fluorescence intensities in the soma were measured and normalized to the average intensity in nontransfected neurons. **C,** Immunofluorescence labeling of untransfected and transfected neurons with GFP-MAP2 for endogenous MAP2. **D,** Schematic diagram showing the domain structure of MAP2C and chimeric proteins with Tau. **E,** Distribution of MAP2C, Tau, and MAP2-Tau chimera. Relative intensity of GFP fluorescence on a line drawn from the base of the axon. **F,** Distribution of GFP-MAP2C and GFP-MAP2C 1SP mutant shown with the immunolabeling of Ankyrin G. Scale bar, 20 µm. **G,** Line scan plots of E. **H,** Quantification of axonal leakage of MAP2C and deletion mutants. See Materials and Methods for details. ANOVA with multiple comparison was used for the statistical analysis (p< 0.0001 for overall difference).

We therefore generated and tested a series of deletion mutants of MAP2C to identify the critical domain. MAP2C primary structure was divided into seven domains based on the amino acid composition and known functions, and individual domains were deleted. Among these deletion mutants, we found that the mutant lacking the S/P domain (ΔSP) exhibited substantial axonal signals unlike WT MAP2c (Fig. 1F and 1G). Mutant lacking the N-terminal half of MAP2c including the S/P domain (Δ1-308) was also distributed to the axon (Fig. 1F – 1H). To test whether the S/P domain is responsible for the change, we put back the S/P domain to Δ1-308. This restored the somatodendritic localization (Fig. 1H), further confirming the importance of the S/P domain in MAP2c localization.

### S/P domain-dependent active transport of MAP2

Stable MT-binding of MAP2 may help keeping MAP2 in the soma and dendrites. To investigate MT-binding of MAP2 in living neurons, we performed fluorescence recovery after photobleaching (FRAP) experiments. Interestingly, the recovery in the second timescale indicated that MAP2 MT-binding is dynamic and that a fraction of MAP2 is in the diffusible state at all times (Fig. 2A). This suggests that there must be mechanisms to keep free MAP2 in dendrites actively. Since MAP2 has been shown to bind to motor proteins in neurons (Gumy et al., 2017), we hypothesized that active transport mechanisms maintain the dendritic localization of MAP2. To test this, we attempted to visualize MAP2 transport in dendrites. We prepared a construct encoding MAP2- tagged with a photoconvertible fluorescent protein, Dendra2 and performed pulse-chase imaging experiments (Fig. 2B). These experiments revealed that photoconverted MAP2 in a small spot in a dendrite spread slowly over time to both proximal and distal directions (Fig. 2C), presumably due to the cycle between diffusion and MT-binding. However, it was clear in kymographs that the redistribution of Dendra2-MAP2 was biased toward distal direction (Fig. 2D), which was also revealed as the shift of line profiles (Fig. 2E). To quantify this, the first frame after photoconversion was subtracted from the frame at 10 min, and signals to proximal and distal sides from the conversion spot were measured (Fig. 2F). In most cases, the signal was redistributed more to the distal direction than the proximal direction of the dendrite (Fig. 2G), indicating slow dendritic transport of MAP2 toward distal dendrites.

**Figure 2.**
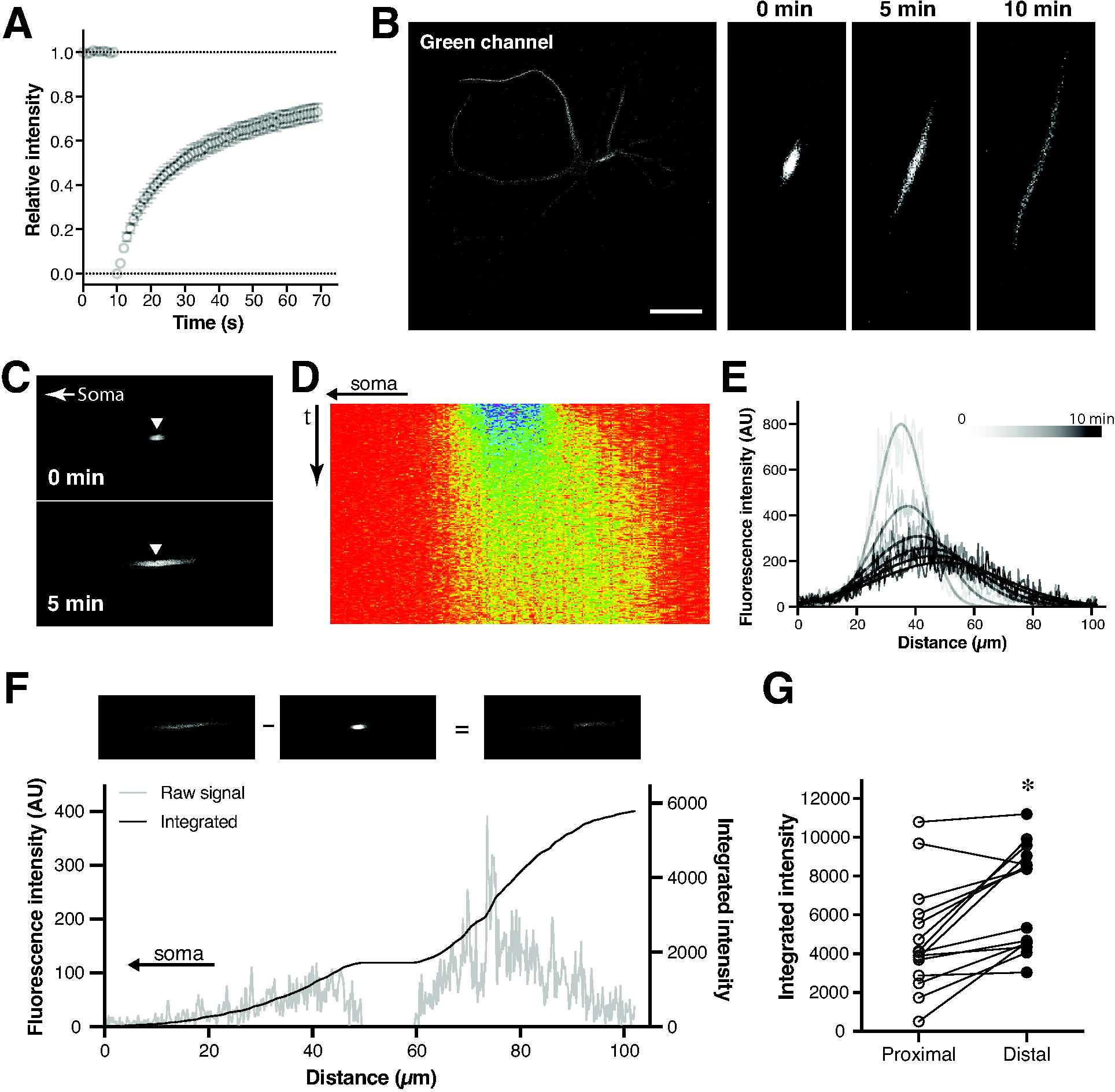
Transport of MAP2C in dendrites. A, FRAP of GFP-MAP2C in dendrites. **B,** Photoconversion of Dendra2-MAP2C in a dendritic branch indicated in the left panel showing the overall distribution before photoconversion. Right panels show the time evolution of red fluorescence after photoconversion. Scale bar, 50 µm. **C,** Migration of photoconverted Dendra2-MAP2C in a dendrite toward the dendritic tip. **D,** Kymograph of C. **E,** Temporal changes of fluorescence distribution over line drawn over the photoconversion spot. **F,** Quantification of the biased distribution of fluorescence signals at 10 min after photoconversion. **G,** Comparison of integrated fluorescence intensities in the proximal and distal sites of the photoconversion spot. *p< 0.0001 using two-way repeated measures ANOVA with multiple comparisons. The analysis was performed with the data of Figs. 3 and 4 to reduce the number of comparisons.

We then investigated whether this dendritic transport is altered by S/P-deletion, which resulted in mislocalization to the axon (Fig. 1). Kymograph analysis after photoconversion in dendrites showed that ΔSP mutant evenly spread from the conversion spot to both proximal and distal directions (Fig. 3A). Quantitative analysis confirmed this (Fig. 3B). The group data also showed that ΔSP no longer exhibited the biased redistribution observed with WT (Fig. 3C). One plausible explanation for this is that more ΔSP is in the free and diffusible fraction than WT. FRAP analysis suggests that this is not the case. ΔSP and WT were virtually identical in the rate and magnitude of fluorescence recovery (Fig. 3D). Taken together, these results suggest that the dendritic localization and dendritic transport both depend on the S/P domain of MAP2.

**Figure 3.**
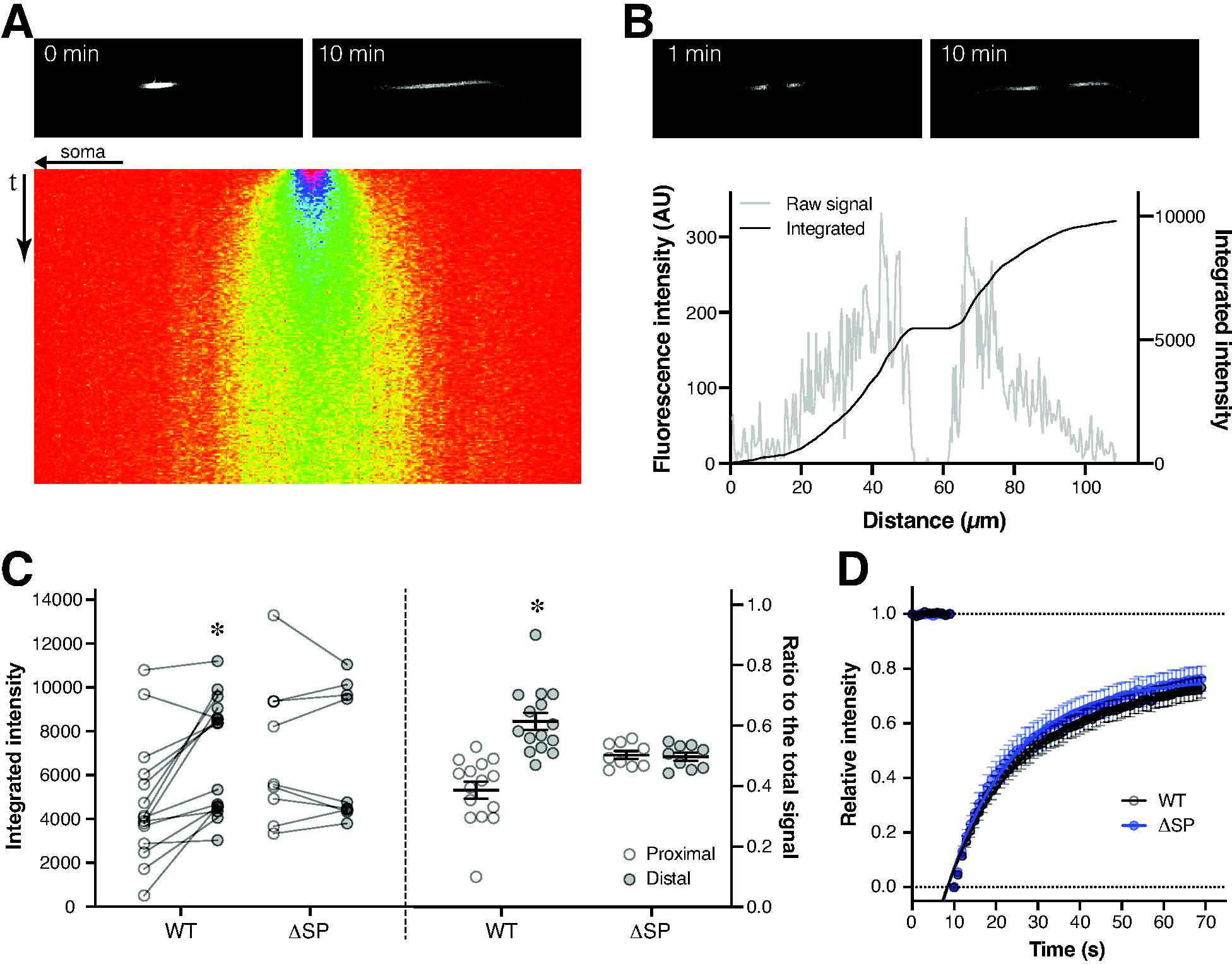
Dendritic MAP2 transport dependent on the S/P region. **A,** Images of Dendra2-MAP2C 1SP immediately after photoconversion and 10 min later (top panels) and corresponding kymograph (bottom). **B,** Quantification of the distribution of fluorescence signals at 10 min after photoconversion. **C,** Comparison of integrated fluorescence intensities in the proximal and distal sites of the photoconversion spot (left) and ratios of intensities over total signal intensities (right). Note that the wild-type (WT) data in the left panel are the same data shown in Fig. 2. There was no significant difference between the proximal and distal values for 1SP (p> 0.99). **D,** FRAP of GFP-MAP2C 1SP in dendrites. WT data from the previous figure are also shown for comparison. The two data sets were fitted with exponential curves with different slopes but could also be fitted with a single curve with no statistical differences.

### Cytoplasmic Dynein-dependent maintenance of MAP2 localization

The slow migration of MAP2 indicates an involvement of motor proteins. What could be the motor protein for the anterograde transport of MAP2 in dendrites? Unlike in the axon, MTs in dendrites have shown to be mixed in their orientations (Baas et al., 1988). If this is the case, motors would not be able to transport MAP2 more preferentially to distal dendrites. However, recent studies suggest that acetylated MTs are oriented uniformly to their minus ends toward the distal end in dendrites (Tas et al., 2017). Since cytoplasmic Dynein transport cargos toward the minus end of MT and binds preferably to acetylated MTs like kinesins (Reed et al., 2006; Dompierre et al., 2007), we considered it as the primary candidate for MAP2 transport. To test this, neurons expressing Dendra2-MAP2 were treated with 50 µM ciliobrevin D, a specific inhibitor of cytoplasmic Dynein (Firestone et al., 2012), for 6 h, and subjected to pulse-chase imaging in the presence of the inhibitor. Intriguingly, the inhibitor disrupted the anterograde transport of MAP2 in dendrites (Fig. 4A), such that there was no obvious biased redistribution of converted MAP2 (Fig. 4B). FRAP analysis showed that ciliobrevin D decreased, but not increased, the recovery slightly but significantly (Fig. 4C).

**Figure 4.**
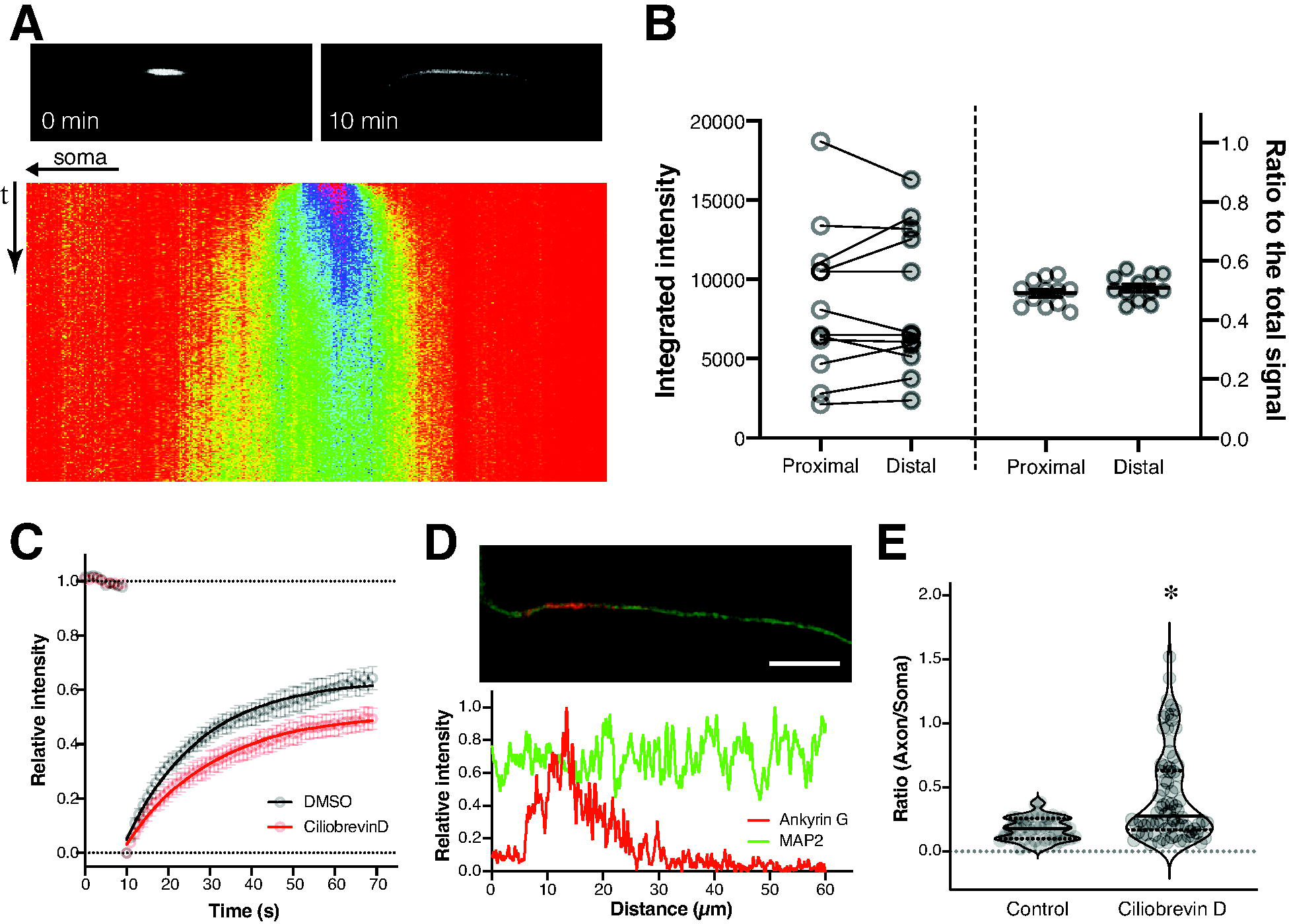
Effects of a Dynein inhibitor, ciliobrevin D, on the dynamics and localization of MAP2. **A,** Images of Dendra2-MAP2C immediately after photoconversion and 10 min later (top panels) and corresponding kymograph (bottom) in neurons treated with 50 µM ciliobrevin D for 6 h. **B,** Comparison of integrated fluorescence intensities in the proximal and distal sites of the photoconversion spot (left) and ratios of intensities over total signal intensities (right). There was no significant difference between the proximal and distal values (p> 0.95). **C,** FRAP of GFP-MAP2C in DMSO-treated control neurons and ciliobrevin D-treated cells. The two data sets were fitted with exponential curves with different slopes but could also be fitted with a single curve with no statistical differences. **D,** Distribution of endogenous MAP2 in neurons treated with ciliobrevin D for 24 h shown by immunofluorescence labeling of MAP2 with Ankyrin G (top). The bottom panel shows the signal intensities of MAP2 and Ankyrin G over a line drawn from the base of the axon. **E,** Ratios of axonal signals over somatic signals of MAP2 immunolabeling in neurons treated with DMSO (Control) or ciliobrevin D. Each data point and violin plots are shown. *p< 0.0001.

We then asked whether the Dynein- and S/P domain-dependent transport is important for the somatodendritic localization of MAP2. To test this, neurons at 14 DIV were treated with 50 µM ciliobrevin D for either 24 h or 48 h and analyzed for endogenous MAP2 localization. Since we obtained more obvious results with the 48 h treatment, we focused on this time point. As shown in Fig. 4D, some of treated cells exhibited clear axonal MAP2 immunostaining in the axon, which was not observed in the DMSO control. The line scan analyses also showed the axonal distribution of MAP2 after ciliobrevin treatment (Fig. 4D). We then quantified the ratio of MAP2 in distal axon over MAP2 signal in proximal axon (hillock) and found that the ratio is significantly greater than control (Fig. 4E). To further substantiate the role of Dynein, we also tested cytoplasmic Dynein knockdown. We used a shRNA for rat cytoplasmic Dynein, which has been reported to reduce it in cultured rat Sertoli cells (Wen et al., 2018) (Fig. 5A). Dynein knockdown did result in the similar leakage of MAP2 into the axon like the pharmacological inhibition (Fig. 5B and 5C). These results suggest that the cytoplasmic Dynein-dependent transport of MAP2 toward distal dendrites is important for the maintenance of the somatodendritic localization of MAP2.

**Figure 5.**
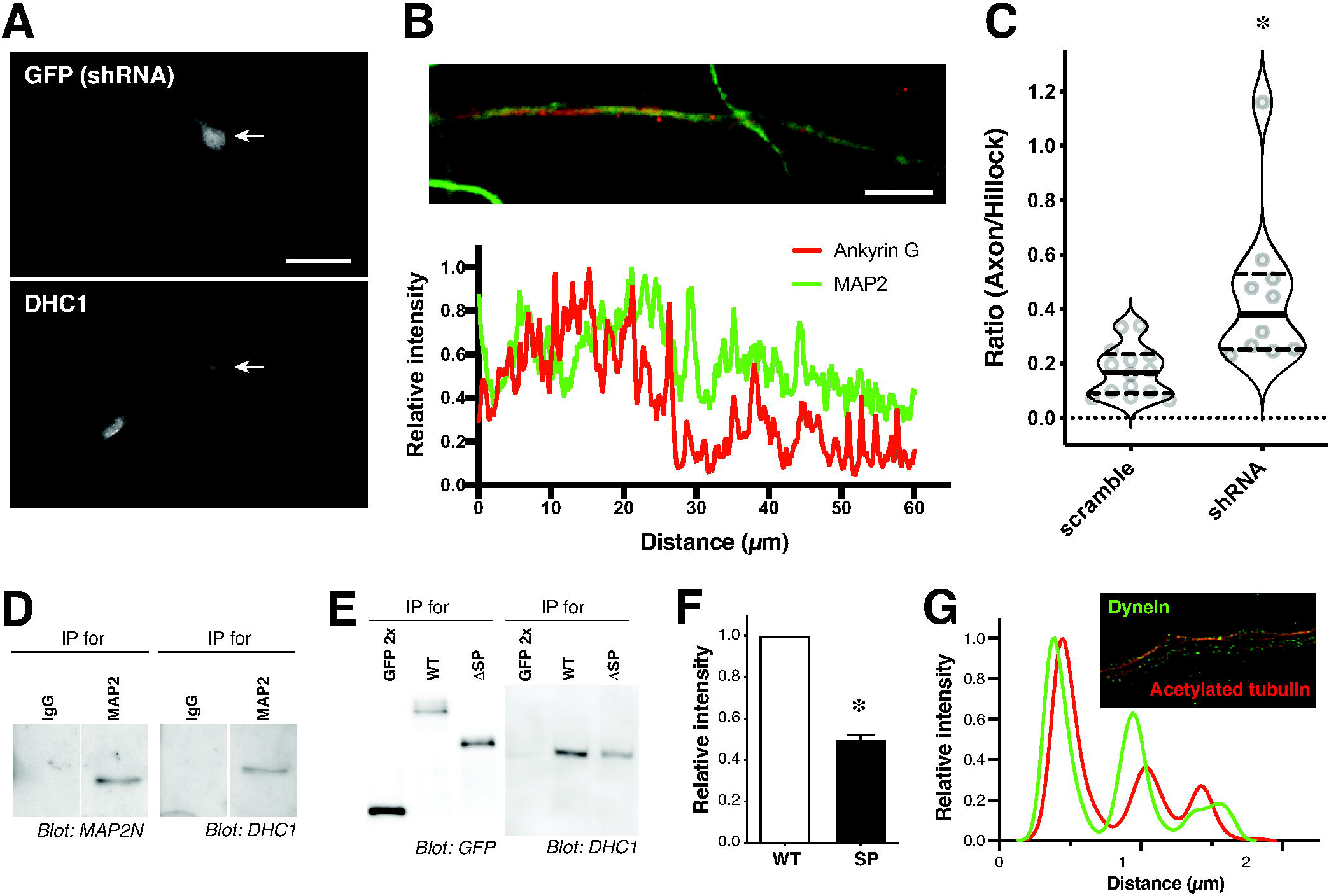
Effects of Dynein knockdown and MAP2-Dynein binding. **A,** DHC1 immunolabeling of neurons transfected with Dynein shRNA plasmid expressing GFP. Scale bar, 50 µm. **B,** Distribution of endogenous MAP2 and Ankyrin G in neurons transfected with Dynein shRNA. MAP2 is shown in a pseudo-color. Scale bar, 20 µm. **C,** Ratios of distal axonal signals over proximal signals of MAP2 immunolabeling in neurons transfected with scrambled or Dynein shRNA. Each data point and violin plots are shown. *p= 0.0002 with Mann-Whitney test. **D,** Immunoprecipitation of MAP2 and co-immunoprecipitation of DHC1 from mouse cerebral lysates. Rabbit IgG was used as negative control. **E,** Co-immunoprecipitation of DHC1 from HEK293 cells expressing DHC1 and WT GFP-MAP2C, GFP-MAP2C 1SP, or tandem GFP (GFP 2x) using an antibody for GFP. **F,** Quantification of signal intensities of co-immunoprecipitated DHC1 with WT and 1SP. *p< 0.0001 with paired t-test. **G,** STED imaging of immunolabeling of DHC1 and acetylated MTs in dendrites (inset). Line scans of fluorescence labeling are shown in the graph.

### Binding of MAP2 to Dynein complex *in vitro* and *in vivo*

We investigated whether MAP2 binds to Dynein transport complex *in vitro* and *vivo* and whether the binding depends on the S/P domain. We first examined their binding *in vivo* in the mouse brain. MAP2 was immunoprecipitated from the brain lysate, and the precipitate was analyzed for MAP2 and cytoplasmic Dynein heavy chain (DHC1). Long isoforms of MAP2 were immunoprecipitated, and DHC1 was also readily detected in the sample (Fig. 5D), indicating that MAP2 and DHC1 can form a protein complex in the adult mouse brain.

We next investigated if the binding of DHC1 depends on the S/P domain of MAP2. First, to reconstitute their binding, wild-type MAP2C and DHC1 were expressed in HEK293 cells, which express endogenous DHC1 at low levels as well (data not shown), and cell lysates were prepared as described in the Materials and Methods. These lysates were incubated with anti-His tag antibody for GFP-MAP2, which has the His-tag between GFP and MAP2. This resulted in the co-immunoprecipitation of MAP2 and DHC1 and demonstrates their *in vitro* binding (Fig. 5E).

However, ΔSP exhibited decreased levels of co-immunoprecipitated DHC1 (Fig. 5E and 5F). Furthermore, as stated earlier, Dynein may utilize acetylated MTs to transport MAP2 toward distal dendrites. Our supoer-resolution microscopy analysis did show dendritic DHC1 labeling along with acetylated MTs (Fig. 5G). These results suggest that the S/P region is critical for DHC1 binding, MAP2 transport, and localization of MAP2, thereby supporting our hypothesis.

### Data-driven model of MAP2 diffusion, binding, and transport

With these quantitative data in hand, we sought to build a mathematical model of MAP2 transport to test whether this kind of transport mechanism is sufficient for the dendritic retention of MAP2 (Fig. 6A). First, we made an estimation of MAP2 diffusion based on FRAP data. We have extensively used FRAP to analyze MT-binding of Tau, a sibling MAP, and previously reported that Tau lacking the microtubule-binding domain (ΔMTBD) is freely diffusible (Iwata et al., 2019). We simulated the FRAP experiment with GFP and ΔMTBD in a thin cylindrical neurite (in this case axon) using the diffusion equation with varying diffusion coefficient under the VCell environment (Fig. 6B). We estimated best fit values by plotting the reciprocals of sum of squares between the data and simulation results and found 22.5 µm^2^/s for GFP and 12.5 µm^2^/s for ΔMTBD as respective best fit values (Fig. 6C). Since wild-type MAP2C is larger than the mutant Tau, we chose 10 µm^2^/s for Dendra2-MAP2C and GFP-MAP2C. These estimates were made based on the assumption that the FRAP procedure results in the full conversion of GFP to a permanently dark state, such that the recovery solely reflects molecular motions. However, it has been reported that GFP variants can be switched to a reversible dark state (Dickson et al., 1997; Sinnecker et al., 2005; Henderson et al., 2007). We performed FRAP after fixing the cells and observed 5% of recovery within 5 s, which we assumed a fraction of GFP in the reversible dark state. We then performed FRAP simulation, in which 5% of GFP goes into the reversible dark state. Fortunately, this did not affect the estimate, presumably due to rapid diffusion of these species out of the bleached region.

**Figure 6.**
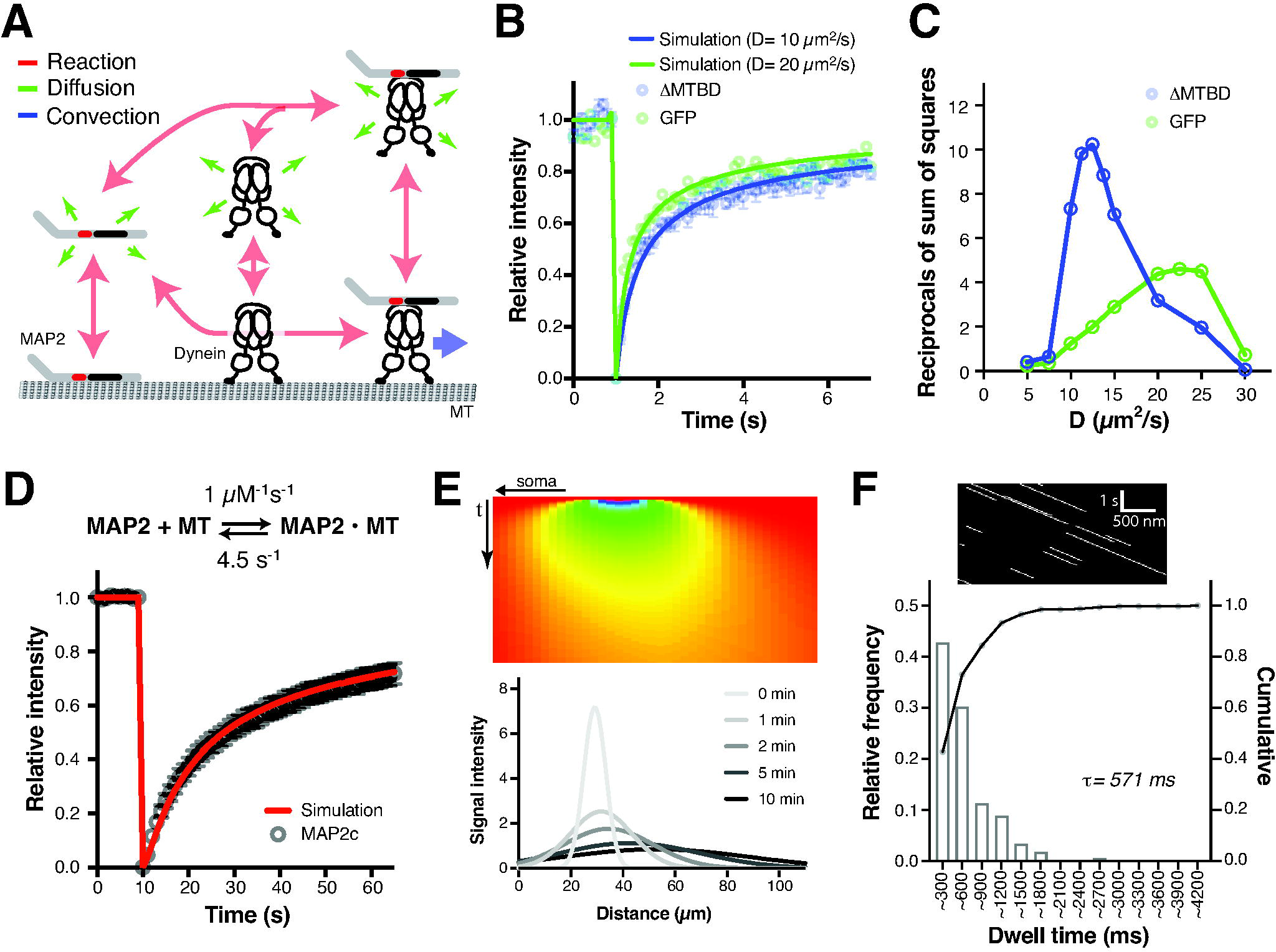
Transport model of MAP2 and parameter estimation. A, Scheme of the transport model. **B,** Estimation of diffusion coefficient from FRAP data of GFP and Tau lacking MTBD (1MTBD) by simulating the FRAP experiment. **C,** Fitting of simulation data on FRAP data with varying diffusion coefficient. Reciprocals of sum of squares are plotted against diffusion coefficient. **D,** Estimation of MT-binding kinetics of MAP2C from FRAP data. The red curve indicates the simulation data with the reaction kinetics shown in the reaction equation. **E,** Virtual kymograph and line scan analysis from model simulation. **F,** Stochastic simulation of the model. The image shows a kymograph of the MAP2-Dynein-MT transport complex, which moves unidirectionally. The graph is the histogram of MAP2 dwell time in the transport complex.

Next, we estimated the concentration of MAP2 in dendrites. The concentration of endogenous MAP2 was estimated from immunoblot quantification of MAP2 in homogenates of adult mouse cortices against purified mouse brain HMW MAP2. Our measurement indicated 1.85 nmol/g wet tissue (ml), which gave about 5.6 µM with an assumption that somata and dendrites occupy one third of the total volume. We employed 5 µM for endogenous MAP2 and 25 µM for GFP-MAP2 because we used neurons expressing 3∼5-fold levels of exogenous MAP2C to endogenous MAP2, based on the results in Fig. 1B, in most of the experiments. We also estimated the concentration of MTs. Based on the structure, there would be about 3,300 tubulin molecules (1,650 α/Δ heterodimers) per 1 µm length of MT. It has been reported that there are on average about 70 MTs/µm^2^ in a dendrite (Kubota et al., 2011; Baas et al., 1988; Katrukha et al., 2021). Therefore, in 1 µm^3^ there could be 230,000 tubulin molecules, which translates to about 400 µM. Assuming that 2∼3 tubulin heterodimers provide a MAP2 binding site with three MT-binding repeats (Kellogg et al., 2018), we estimated 75 µM potential MAP2-binding site (MTs in the simulation). In an immunoblot quantification, we obtained about 80 nmol/g wet tissue of α-tubulin. Therefore, we estimated 320 µM total (α and Δ) tubulins and 55 µM MTs, if neurons occupy 50% of the tissue volume, and if 85% of tubulins contributes to MTs (Hagita et al., 2021). We utilized 75 µM MTs in the following simulations but obtained comparable results with 40 µM MTs with minor adjustments in the binding affinity between MAP2 and MTs (see below).

We then simulated FRAP data of wild-type GFP-MAP2 using the reaction-diffusion equation with varying binding parameters. The data were fitted well with KD= 4.5 µM, when Kon was set above 0.1 µM^-1^s^-1^ (Fig. 6D). We mainly used Kon of 1 µM^-1^s^-1^ for the following simulations, as there were no discernable differences between 0.1 and 1 µM^-1^s^-1^. With 40 µM MTs, the data were fitted well with KD= 1 µM. Diffusion coefficient and MT-binding parameters of Dynein were determined based on literatures. MAP2-Dynein affinity was determined based on the affinity for linker peptides (Lee et al., 2018; Lee et al., 2020). Concentration of Dynein was set at 5 µM arbitrarily. Parameters used for the model are summarized in Table 1.

**Table 1.** Condition for the simulation

| **Diffusion coefficients** |  | **Binding affinity ($K_D$)** |  | **Initial concentrations** |  |
| --- | --- | --- | --- | --- | --- |
| Free MAP2 | 10 $\mu\text{m}^2/\text{s}$ | MAP2-MT | 4.5 $\mu\text{M}^*$ | Endogenous MAP2 | 5 $\mu\text{M}$ (0 in axon) |
| Dynein-bound MAP2 | 1 $\mu\text{m}^2/\text{s}$ | MAP2-Dynein | 10 $\mu\text{M}^{\S}$ | Dendra2-MAP2 (green) | 25 $\mu\text{M}$ (0 in axon) |
| MT-bound species | 0.001 $\mu\text{m}^2/\text{s}$ | Dynein-MT | 2 $\mu\text{M}^{\P}$ | Dendra2-MAP2 (red) | 0 $\mu\text{M}$ |
| | | | | Dynein | 5 $\mu\text{M}$ |
| **Motor speed** | 1 $\mu\text{m}/\text{s}$ | | | MT (binding sites) | 75 $\mu\text{M}$ |

\*The value was estimated based on the initial putative $K_{on} = 1 \mu\text{M}^{-1}\text{s}^{-1}$ . Note that $K_D = 3.3 \mu\text{M}$ with $K_{on} = 0.09 \mu\text{M}^{-1}\text{s}^{-1}$ provided comparable fit to the FRAP data and worked similarly on simulation. $\S K_D$ was estimated from literatures for linker-Dynein binding (Lee et al., 2018; Lee et al., 2020) and from the Co-IP experiment. $K_{on}$ used for this estimation was $0.1 \mu\text{M}^{-1}\text{s}^{-1}$ . $\P$ The $K_D$ was based on previous reports (Mizuno et al., 2004; Gibbons et al., 2005; Ananthanarayanan et al., 2013; Zhang et al., 2017) and $K_{off}$ from Dynein dwell time on MTs (Zhang et al., 2017).

To simulate the dendritic photoconversion imaging, a 200 µm long cylinder with 3 µm diameter was built in the VCell. To implement active transport, reaction-diffusion-convection (advection) equations were applied to Dynein species bound to both MT and MAP2. To avoid redistribution of MTs, which are expected to be virtually immobile, the concentration of MT was made constant in each unit volume. The velocity of convection, i.e., the speed of motor, was set to 1 µm/s. Photoconversion took place in the middle of the cylinder. Virtual kymograph of the converted species (red MAP2) showed a gradual spreading of the species due to diffusion and drifting of the center of distribution because of the transport (Fig. 6E). This was similar to the kymograph from imaging experiments. The line scan profiles were also similar to the data (Fig. 6E). We then tested this model in a bottom-up particle-based stochastic simulation using Smoldyn based on the Smoluchowski model of diffusion influenced systems (Andrews, 2012) in the VCell environment with the same parameters. This enabled us to estimate how individual MAP2 molecules behave with the given condition. The model estimated that about 87% of MAP2 is bound to MTs, 8.4% free, and 4.2% being transported (bound to motor and MT) at a given point in time. With the Kon and Koff we used MT-binding was transient with median and τ of dwell time at 140 and 239 ms, respectively, which is comparable to the predicted residence time (1/Koff), 222 ms. MAP2 in the transport state was also short-lived, such that the dwell time in the transport state was less than 1 s in 90% of the case (Fig. 6F). This would account for the slow and gradual transport of MAP2 in dendrites.

### Testing the capability of the mechanism in localizing MAP2

We then examined whether this intermittent transport mechanism could retain MAP2 in the somatodendritic region without substantial leakage to the axon in simulation, as observed in neurons. Simplified neuron was constructed with a 20 µm diameter spherical cell body, 100 µm long cylindrical dendrites with 3 µm diameter, and a 400 µm long cylindrical axon with 2 µm diameter (Fig. 7A). Convection for the MAP2-Dynein-MT complex was implemented in the axon and dendrites but not in the soma. Because we do not have a return mechanism for Dynein we had to keep the Dynein concentration constant in all compartments and to reduce the speed of convection toward the tips of dendrites (Fig. 7A). Otherwise, the motor and cargo species accumulated at the tips to extremely high concentrations. Analysis of green MAP2 species provided FRAP simulation in this model and revealed similar but slightly faster recovery after bleaching, due to the convection. Therefore, the binding kinetics was slightly modified to fit the data (Kon from 1 µM^-1^s^-1^ to 1.5 µM^-1^s^-1^ and KD from 4.5 to 3 µM).

**Figure 7.**
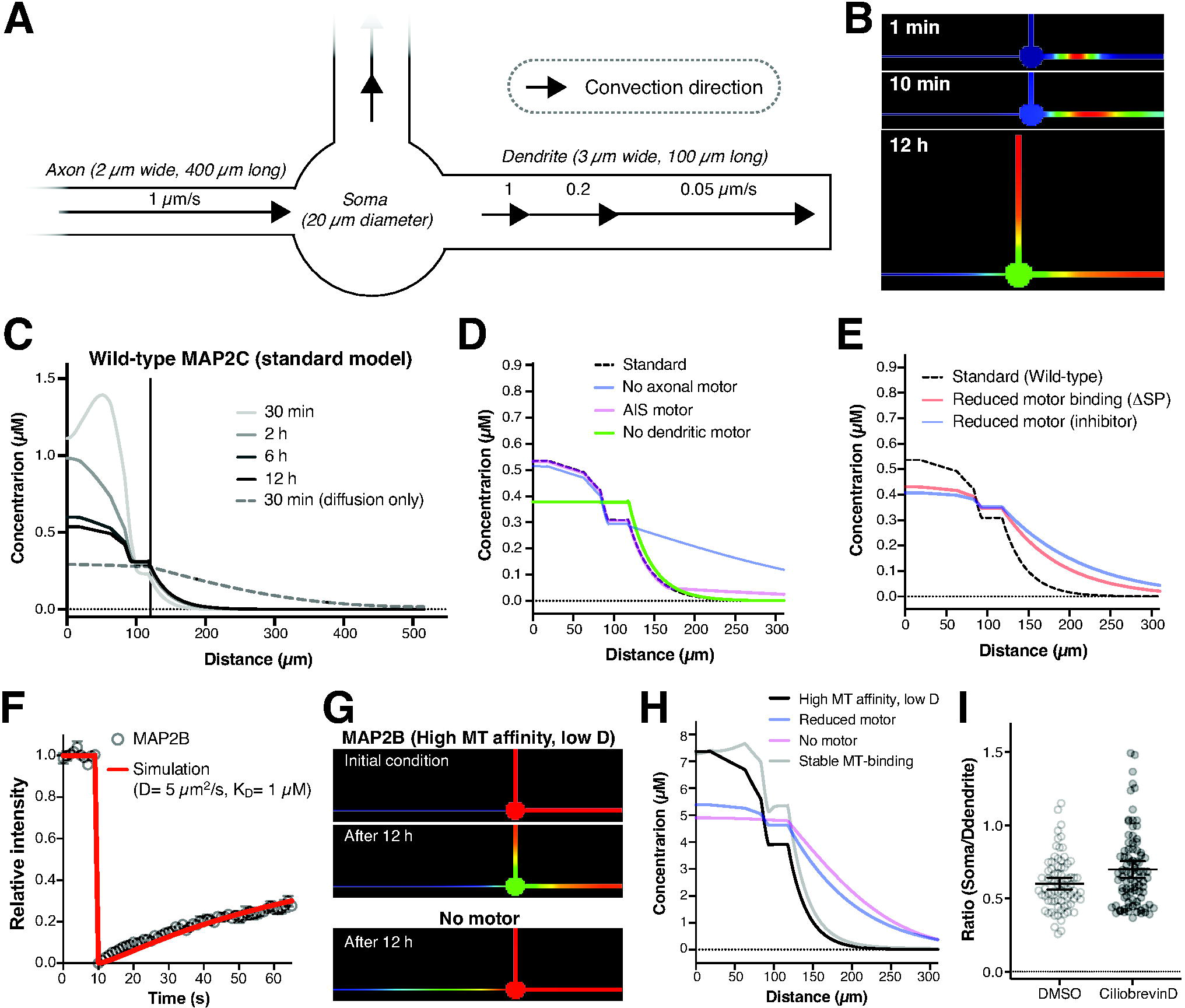
Implementation of the transport model in a virtual neuron. A, Illustration of the virtual neuron geometry and motor activity. **B,** Virtual photoconversion experiment. Concentration of the converted red species is color-coded in a blue-to-red spectrum. **C,** Concentration changes from a dendrite, where photoconversion took place, to the tip of the axon at different time points after conversion. The vertical line indicates the border between the soma and the axon. The dotted curve is for MAP2 lacking MT-binding. **D,** Virtual photoconversion experiments with different motor placement in the neuron. The dotted curve indicates the standard model shown in C. **E,** Virtual photoconversion experiments to mimic S/P deletion and Dynein inhibition. The dotted curve indicates the standard model shown in C. **F,** Estimation of MT-binding kinetics of MAP2B from FRAP data. The red curve indicates the simulation data with the diffusion coefficient and reaction kinetics indicated. **G,** Virtual photoconversion experiment with the MAP2B parameters with or without the motor species. Concentration of the converted red species is color-coded in a blue-to-red spectrum. **H,** Virtual photoconversion experiments of MAP2B with different motor activity and MT-binding in the neuron. **I,** Reduction of the immunofluorescence signal gradient of endogenous MAP2 between dendrites and the soma after ciliobrevin D treatment.

We then followed the distribution of red MAP2 over 12 h in the simulation (Fig. 7B). It was retained in the original dendrite at 30 min after conversion, as observed in experiments (Fig. 2A), but distributed to the soma and the other dendrite over time (Fig. 7C). There was a slight leak into the axon (Fig. 7B and 7C). However, the concentration in the axon decays quickly within the first 50 µm (Fig. 7C). This was also observed for green MAP2 and endogenous MAP2, which were situated only in the soma and dendrites at the beginning (not shown). Almost identical results were obtained with 40 µM MT and KD= 1 µM (not shown). Considering the thinness of axons, this leak would be negligible.

We then tested whether the motor is necessary for the entire length of the axon (Fig. 7D). Motor concentration was homogeneous throughout the model cell. Dendritic motor was found necessary to establish the concentration difference between the soma and dendrites. When we removed the motor from the axon entirely, MAP2 species were distributed to distal axon after 12 h. Then, we put it back only to the first 50 µm segment of the axon (AIS). This dramatically reduced the axonal leak. However, beyond AIS MAP2 still exhibited slight leak into distal axon. Therefore, axonal retrograde transport of MAP2 plays an essential role in the model with the estimated affinity of the motor to MTs and MAP2. Unfortunately, we were not able to perform the photoconversion experiments in the axon, as the levels of Dendra2-MAP2 were not sufficient in the axon. Since Dynein is known to distribute throughout the axon and contribute to the retrograde axonal transport, we implemented the motor-based transport in the entire length of the axon.

Intriguingly, we could replicate the substantial axonal leak of MAP2 with the deletion of the S/P domain and ciliobrevin by reducing either the binding affinity between MAP2 and motor (from KD= 10 µM to 30 µM) or the concentration of active motor (from 5 to 1.5 µM) (Fig. 7E). However, the experiments with ciliobrevin were done on endogenous MAP2, most of which is likely to be MAP2A and MAP2B. Therefore, we investigated whether these large isoforms can be localized by the same mechanism. FRAP analyses of GFP-MAP2B exhibited much slower recovery than MAP2C (Fig. 7F). To fit to the data, diffusion coefficient and KD were reduced to 5 µm^2^/s and 1 µM, respectively (Fig. 7F). The whole cell simulation was performed with these new parameters and revealed that MAP2B behavior is quite similar to that of MAP2C (Fig. 7G). Also, reducing the concentration of motor to mimic the effect of ciliobrevin and shRNA, as well as removing it entirely, resulted in a significant leak of MAP2B into the axon (Fig. 7H). Moreover, simulation results predicted that Dynein inhibition results in not only the axonal leak but also dissipation of the somatodendritic concentration gradient. We therefore reanalyzed the immunostaining data and found that indeed the somatic signals were increased in ciliobrevin D-treated cells (Fig. 7I). These results therefore indicate that our transport model captures the somatodendritic retention of MAP2 and suggest that the motor-dependent mechanism is critical for MAP2 localization.

### Dissecting the roles of MT-binding, diffusion barrier, and local metabolism

There have been three canonical models for the somatodendritic localization of MAP2 beside our model: Preferential binding model, AIS restricted diffusion model, and local degradation model. We therefore investigated the potential roles of these mechanisms in MAP2 localization by modeling them. It has been proposed that MAP2 is tightly bound to dendritic MTs, more so than to axonal MTs, and therefore retained in the compartment (Fig. 8A). This may not be likely because FRAP of GFP-MAP2C in dendrites and the axons, in which trace levels of GFP-MAP2C was detectable, were virtually identical (Fig. 8B). Nevertheless, to test this model, we implemented the data-based MT-binding in dendrites, while putting reduced MT-binding in the soma and axon. This resulted in a substantial leak into the axon in the absence of the motor (Fig. 8A), suggesting that the preferential binding model is insufficient by itself.

**Figure 8.**
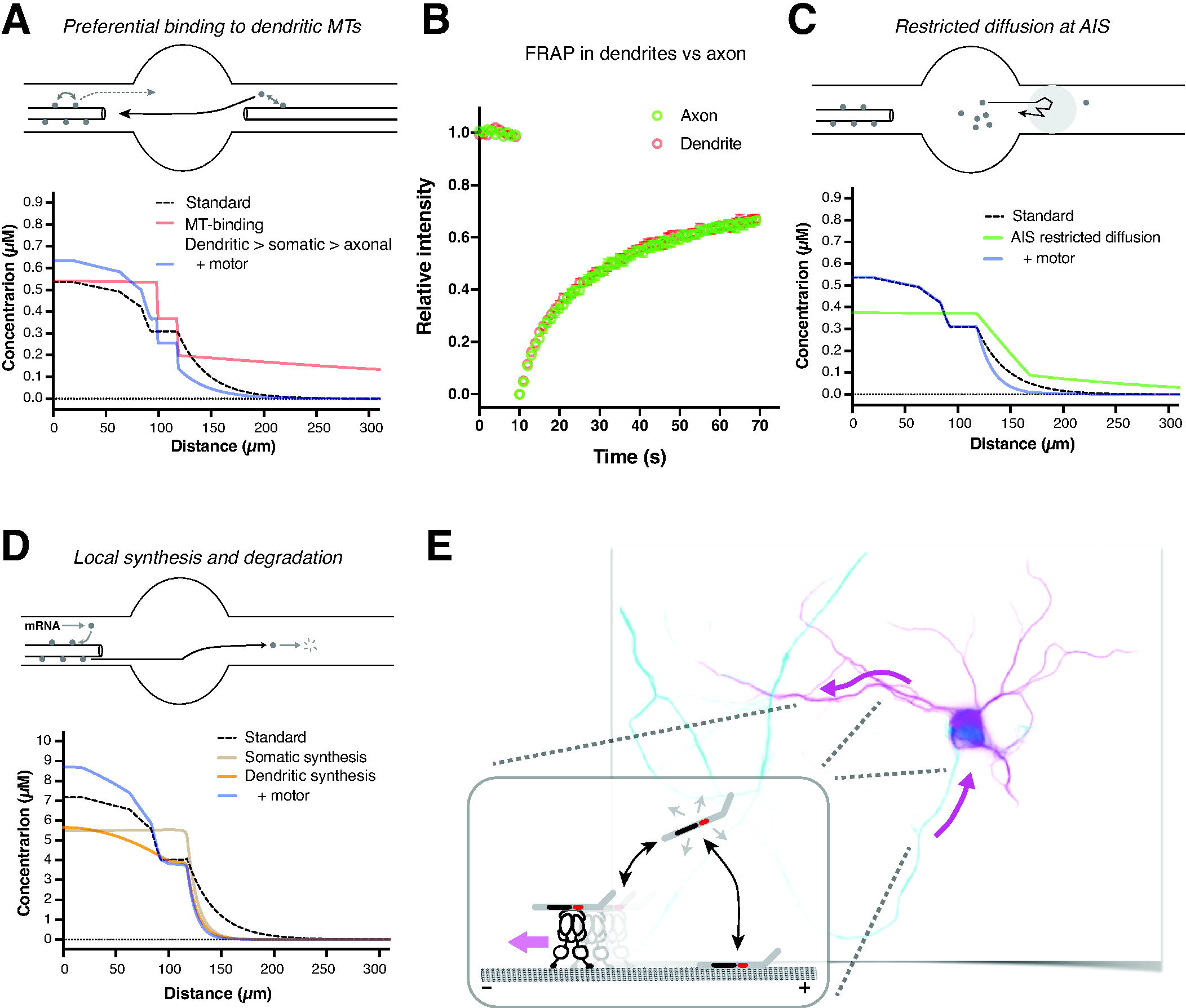
Modeling the other proposed mechanisms of MAP2 localization. A, Preferential binding of MAP2 to dendritic MTs as illustrated in the top panel. Virtual photoconversion experiment was performed with compartment specific KD to MTs (3 µM in dendrites, 4.5 µM in the soma, and 9 µM in the axon) without or with motor. The dotted curve indicates the standard model. B, FRAP of GFP-MAP2C in a dendrite and the axon. C, AIS restricted diffusion model. Virtual photoconversion experiment was performed with compartment specific diffusion coefficient (1 µm^2^/s in the first 50 µm segment and 10 µm^2^/s for the rest) without or with motor. D, Local synthesis and degradation model. New seed species mimicking mRNA and dead species mimicking degradation were added to the model. The synthesis reaction had the forward reaction rate of 0.0001 s^-1^, and the degradation rate was 0.1 s^-1^. The synthesis reaction was implemented to the soma or dendrites. E, Illustration depicting our transport model of MAP2 for the somatodendritic localization.

Another canonical hypothesis is that MAP2 entrance to the axon is restricted at the AIS by either a hypothetical diffusion barrier or restricted diffusion (Fig. 8C). To test this model, the diffusion coefficient of MAP2 in the first 50 µm segment of the axon was reduced from 10 µm^2^/s to 1 µm^2^/s. This “AIS diffusion barrier” was not sufficient to prevent the axonal leakage of MAP2 in the absence of motor (Fig. 8C). Also, it did not reproduce the concentration gradient between the soma and dendrites (Fig. 8C). Therefore, the AIS restriction model is also insufficient to account for the somatodendritic localization of MAP2.

Lastly, we tested the local degradation model, based on that MAP2 is degraded within the axon in spinal neurons (Fig. 8D). Since this model naturally requires balanced synthesis of MAP2, we introduced somatic synthesis of MAP2. This failed because it did not create the soma-dendrite gradient at all (Fig. 8D). So, we eliminated the somatic synthesis and introduced local synthesis of MAP2 in dendrites instead. The combined model exhibited very small axonal leakage (Fig. 8D). However, the concentration gradient between the soma and dendrites was still very small (Fig. 8D). Together with that cDNA of GFP-MAP2 does not contain any regulatory untranslated regions, which are typically required for local translation of mRNA, yet localize properly and that Dynein inhibition changed the localization, this mechanism is considered not to be essential.

Even though the aforementioned models were found to be insufficient, they may refine the localization of MAP2 established primarily by the active transport mechanism. We introduced the motor species into these models and found a substantial improvement of the soma-dendrite concentration gradient and/or axonal leakage (Fig. 8A, 8C, and 8D). Taken together, these experimental and theoretical results suggest that the Dynein-based active transport mechanism is the key for the somatodendritic localization of MAP2 potentially supported by the other mechanisms.

## DISCUSSION

MAP2 has been widely and extensively used as the marker of dendrites. In addition to its predicted role in stabilizing MTs in dendrites (Itoh and Hotani, 1994), MAP2 has recently been shown to play a role in cargo selection in the junction between the soma and AIS (Gumy et al., 2017). Despite the importance of MAP2 localization in research and physiology, the mechanism for its specific and discrete somatodendritic localization has been elusive. In this study, we tackled this longstanding question of neurobiology using quantitative imaging and computational approaches. Particularly, we developed a method to estimate diffusion coefficient and MT-binding kinetics constants of MAP2 from FRAP data by fitting with virtual FRAP data from numerical simulation. With the pulse-chase imaging data and the data-based molecular parameters of MAP2, we constructed a mathematical model of motor-driven MAP2 transport and demonstrated that this rather simple mechanism is sufficient to establish and maintain the somatodendritic localization of MAP2 in simulation.

Using a series of deletion mutants, we found that somatodendritic localization of MAP2 is regulated and maintained in a S/P-dependent manner. Using the pulse-chase imaging approach, we also found that MAP2 movement is biased to the tip of dendrites and this biased migration is lost when cells are treated with a Dynein inhibitor, Ciliobrevin D, or when the S/P region is removed from MAP2. In addition, we demonstrated that MAP2 interacts with DHC1 via the S/P region. These results suggest that Dynein complex transports and retains MAP2 in dendrites, resulting in somatodendritic localization of MAP2. This Dynein-dependent active transport mechanism (Fig. 8E) is further supported by numerical simulation constrained by data-driven parameters.

Our imaging experiments rely on fluorescent protein-tagged MAP2 overexpressed in cultured neurons. This may raise a concern that observed phenomena are specific to these species but not to the endogenous proteins. Fortunately, we were able to show that Dynein inhibition and knockdown disrupted the localization of endogenous MAP2, and that endogenous MAP2 in the mouse brain can also interact with Dynein. Based on these findings together with imaging data and simulation results, we conclude that transport of MAP2 via its binding to Dynein is essential to establish and maintain the somatodendritic localization in neurons.

### Dynein-based transport mechanism

Our transport model stems on the experimental results that transport and localization of MAP2 are inhibited by S/P deletion and Dynein inhibitor or knockdown, and that Dynein coimmunoprecipitates with MAP2, which is also inhibited by S/P deletion. Therefore, we hypothesize that Dynein complex captures free MAP2 and maintains MAP2 in dendrites via its transport activity. One aspect we have not been able to demonstrate experimentally was retrograde transport of MAP2 in the axon, due to insufficient levels of Dendra2-MAP2 present in the axon to perform the photoconversion experiments. Our simulation analyses clearly showed that MAP2 leaks into the axon substantially without the axonal retrograde transport, indicating that axonal transport toward the soma is an essential part of the localization mechanism, at least in the absence of somatic retention mechanism we did not implement.

As stated above, the model is supported by the results that MAP2 binds the Dynein complex via the S/P region. However, one might argue that because both proteins bind to MTs they can coimmunoprecipitate through MT-binding, instead of endogenous Dynein light and intermediate chains or adaptor proteins, some of which are expressed in HEK293 cells (Sachdev et al., 2007). In this scenario, Dynein would not be able to transport MAP2 directly. In our co-IP experiment using HEK cells, lysate was prepared after more than 40 min long cold treatment on ice or at 4℃. This long cold treatment is known to induce MT depolymerization. Therefore, it is unlikely that sufficient MTs survived in our preparations. Also, FRAP experiments did not show any substantial reduction of MT-binding of ΔSP mutant, which would be expected if the coimmunoprecipitation relies on MT-binding of MAP2 and Dynein. Therefore, we conclude that the interaction of MAP2 with DHC1 is not mediated by MTs.

The S/P region of MAP2 consists of 15% proline and ∼20% serine. Does this characteristic amino acid composition, which might result in specific conformation such as poly proline ii helix (Melková et al., 2018), need to be recognized by Dynein transport system? As previously reported, the proline rich region of Herpes virus viral protein 1/2 (VP1/2, also called pUL36) has been shown to interact with Dynactin components, a part of the Dynein motor complex, resulting in enhanced transport to the soma by Dynein (Zaichick et al., 2013). Unfortunately, there is no obvious similarity between the S/P region of MAP2 and the proline rich region of VP1/2. Also, we did not find any similar sequences to that of S/P in the human genomic database. Identifying the actual Dynein-binding motif in the S/P region and similar sequences in other proteins might allow us to generalize the transport model to localizations of other dendritic cytoplasmic proteins.

It has been reported that mammalian Dynein does not move processively along MTs on its own in *in vitro* assays (McKenney et al., 2014), unlike yeast Dynein (Reck-Peterson et al., 2006). Dynein needs cofactors, Dynactin (p150 Glued) and cargo adaptors, BicD2, Rab11-FIP3, Spindly or Hook3 to move processively along MTs (McKenney et al., 2014). Dynactin (p150 Glued) is reported to bind to tyrosinated tubulin more preferentially than to detyrosinated tubulin, whereas DHC1 does not show such preference (McKenney et al., 2016). This binding to tyrosinated MTs appeared to aid the initiation of Dynein processivity, although the processivity itself does not require tyrosinated MTs. The orientation of tyrosinated tubulin is reported to be plus-end out in the soma and dendrites (Tas et al., 2017). Our imaging data showed that MAP2 is transported to the tip of a dendrite and that this biased transport was canceled by Dynein inhibitor, suggesting that transport is mediated by Dynein along acetylated MT that is oriented minus-end out in dendrites. This seems contradictory to the previous reports. We speculate that the Dynein complex carrying MAP2 may contain different Dynein cofactors, which prefer acetylated MTs, or that acetylated MTs are not completely devoid of tyrosination.

### Dynamic MT-binding

Our FRAP data and stochastic simulation suggest that MT-binding of MAP2 in living neurons is quite dynamic with estimated KD of 1∼3 µM (depending on the estimated level of MTs). This was surprising because MAP2 binds to MTs quite stably in *in vitro* binding assays (Itoh et al., 1997). Overexpressed GFP-MAP2 might saturate MTs, thereby increasing a fraction of free GFP-MAP2. Our empirical and experimental estimation show that there is 60∼100 times more tubulin molecules than MAP2 in the cytoplasm. Therefore, MAP2 binding sites (MTs) are likely to be abundant enough to capture the estimated level of expressed MAP2. A recent paper using super-resolution microscopy indicates that there could be even more MTs in dendrites than our estimate (Katrukha et al., 2021). This may be the reason why we did not see any changes in endogenous MAP2 despite the five-fold overexpression of GFP-MAP2C, and that overexpressed MAP2C was retained in the somatodendritic region despite co-overexpression of Tau (see Fig. 1A).

In our previous study on the Tau protein, we showed that most of overexpressed Tau could bind to MTs and therefore exhibited very slow recovery in FRAP, when the phosphorylation sites in the proline-rich region of Tau was mutated to alanine to mimic dephosphorylation (Iwata et al., 2019). We also found that MT-binding of wild-type Tau is quite dynamic like that of MAP2 with faster recovery than the alanine mutant, and that pseudo-phosphorylation mutant with glutamate instead of alanine substitutions exhibited recovery even faster than wild-type Tau (Iwata et al., 2019). These results indicated that wild-type Tau is phosphorylated at certain residues in the proline-rich region and thus exhibits dynamic MT-binding in neurons. Based on these findings, we speculate that MAP2 is also phosphorylated at certain residues in neurons (Illenberger et al., 1996; Jansen et al., 2017), and that the total lack of phosphorylation in *in vitro* assays results in the difference observed (Illenberger et al., 1996).

Is the dynamic MT-binding beneficial? For MAP2 localization, it may be. We mainly used KD of 3 µM with Kon of 1.5 µM^-1^s^-1^, which may be considered a relatively fast reaction. We also tested lower Kon at 0.1 µM^-1^s^-1^, which worked as well on FRAP data fitting and the transport simulation. However, when the KD of MAP2 to MTs was decreased from 3 µM to 0.4 µM with the slow reaction kinetics (Kon= 0.1 µM^-1^s^-1^), which was derived from the pseudo-dephosphorylation mutant of Tau (Iwata et al., 2019), in simulation, MAP2 was still distributed mainly to dendrites but with substantially higher levels of axonal MAP2 (Fig. 7H). This is due to that MAP2 strayed into the axon lingers longer due to the stable binding to MTs. Therefore, MT-binding of MAP2 needs to be regulated, presumably by phosphorylation, and the resultant dynamic binding of MAP2 to MTs results in efficient transport and somatodendritic localization. Another consideration is that longer residence of MAP2 on MTs may interfere with motor processivity. MAP2 has been shown to block the path of kinesin motors in *in vitro* assays (von Massow et al., 1989; Hagiwara et al., 1994; Lopez and Sheetz, 1993; Seitz et al., 2002; Monroy et al., 2020). If this is also the case in neurons, lingering MAP2 on MTs may similarly inhibit motor-based transport and possibly its own transport. We are currently not able to implement fiber representation of MTs in the simulation. Further studies are needed to address this issue.

### Other models for the somatodendritic localization of localization of MAP2

As explanations for the somatodendritic localization of MAP2, there have been essentially four models: PD interfering entering of MAP2 into axon (Kanai and Hirokawa, 1995), AIS diffusion barrier preventing MAP2 entrance, preferential binding to dendritic MTs (Kanai and Hirokawa, 1995), and local translation in dendrites with axonal degradation (Garner et al., 1988) (Hirokawa et al., 1996). We cannot fully exclude the possibility that previously reported PD affects localization of MAP2. However, we showed that MAP2C natively lacking PD still exhibited somatodendritic localization, indicating that PD is not necessary for the somatodendritic localization of MAP2.

Diffusion of membrane proteins are clearly restricted in the AIS (Winckler et al., 1999; Nakada et al., 2003), presumably due to crowding in the membrane. Diffusion barrier for cytoplasmic proteins like MAP2 is much less known, and the corresponding physical entity is difficult to comprehend. For MAPs, reinforced MT-binding in the AIS could make overall diffusion slower. We performed FRAP in the first 50 µm of the axon to see if we could detect difference in recovery. There were no differences (See Fig. 8B). Nevertheless, some diffusion restriction of cytoplasmic proteins including fluorescent proteins has been reported (Song et al., 2009). We therefore performed simulation of a model in which MAP2 diffusion is slowed down ten-fold. This diffusion barrier did limit axonal leakage of MAP2 but not sufficiently without the motor-driven transport.

Regarding preferential binding of MAP2 to dendritic MTs, our FRAP experiments showed no difference in recovery in the axon and dendrites, suggesting that MT-binding is comparable in these two compartments. Although some posttranslational modifications such as acetylation differ between the axon and dendrites in very immature neurons (Hammond et al., 2010), they are more uniformly present in in both dendrites and the axon in neurons after 7 DIV (unpublished observations), which we used in the study. Furthermore, MAP2B, which appeared to bind to MTs more stably than MAP2C does, still mislocalized to the axon, when the motor is inhibited, both in the experiments and simulation. These findings indicate that the preferential binding model is insufficient to account for the somatodendritic localization of MAP2.

In contrast, the local synthesis and degradation model may be sufficient for MAP2 localization, when they are combined, as the simulation indicated. However, we (this paper) and others have shown that expressed MAP2 without any native mRNA structures such as a 3’- untranslated region, which are typically necessary for mRNA transport and local translation (Fernandopulle et al., 2021), still localizes properly to dendrites (Kaech et al., 1996). Therefore, local translation of MAP2 in dendrites may not be necessary. Then, can local degradation of MAP2 in the axon alone be sufficient? We also run the simulation of axonal degradation model with the conventional somatic synthesis of MAP2 without dendritic synthesis. While it did restrict the axonal leakage of MAP2 very well, it could not create the concentration gradient between dendrites and the soma. Therefore, for this model to work, dendritic synthesis and axonal degradation must be combined and precisely balanced.

In summary, our findings suggest that active transport of both LMW and HMW MAP2 by Dynein is a key to maintain somatodendritic specific localization of MAP2.

## Materials and Methods

### Neuronal culture

Dissociated cultures of embryonic (E17∼18) hippocampal neurons were prepared from female timed pregnant Sprague Dawley rats as previously described (Misonou et al., 2008) with minor modifications. Briefly, dissected hippocampi were digested in 0.25% trypsin for 15 min at 37°C, dissociated by pipetting, and then plated onto glass coverslips coated with 1 mg/ml poly-L-lysine at 50,000 cells/coverslip. Neurons were cultured for 3 to 24 days *in vitro* (DIV) in 6-well plates, on the bottom of which contained astrocyte cultures. Coverslips were lifted with wax pedestals as described by Kaech and Banker (Kaech and Banker, 2006). Cytosine arabinoflanoside was added to the culture at 2 DIV to prevent the growth of non-neuronal cells on the coverslips. All animal use was approved by the institutional animal care and use committee.

### DNA plasmids

Human MAP2C cDNA tagged with 6xHis was inserted into pAcGFP (Takara) vector using In- Fusion cloning kit (Takara). MAP2C ΔSP was created by removing “S/P region” (P134-P251 in the numbering of 471 amino acids isoform, MAP2C) from pAcGFP-His-MAP2C by PCR. MAP2C Δ1-308 was created by removing 1-308 amino acids from pAcGFP-His-MAP2C by PCR. MAP2C Δ1-308 + SP was created by amplifying the S/P region, which was inserted into pAcGFP-His- MAP2C Δ1-308 by In-Fusion cloning kit (Takara). pAcGFP-His-MAP2C-Tau was chimera of MAP2C_M1-L311 and Tau0N4R_P193-L383 in the numbering Tau0N4R, which was created by PCR and the chimera MAP2C-Tau was amplified by PCR and inserted into pAcGFP vector using In-Fusion cloning kit. Dendra2-MAP2C was prepared as following. pAcGFP-His-MAP2C without AcGFP was amplified and Dendra2 was amplified from pDendra2-C vector (Takara). Dendra2 was inserted into where AcGFP was by In-Fusion cloning kit. Dendra2-MAP2C ΔSP was prepared by removing the S/P region from Dendra2-MAP2C by PCR. MAP2B cDNA was amplified from human hippocampus cDNA library and inserted into where MAP2C was in pAcGFP-His-MAP2C. To express shRNA targeting rat DHC1, rat DHC1-targeting oligonucleotides, AATTCAAGCTGGCGTGCAA, was inserted into BglII and HindIII site of pSUPER-retro-neo-GFP vector according to manufacture manual. All constructs were sequenced to confirm the sequences. DHC1 plasmid was kindly gifted by Dr. Yoko Toyoshima (The University of Tokyo) and we inserted Flag tag into streptavidin binding peptide tag, which locates in the N-terminal region of the plasmid (or just inserted Flag tag into N-terminus of the plasmid).

### Transfection

Neurons were transiently transfected at 7 DIV using Lipofectamine 2000 transfection reagent (Thermo Fischer Scientific, Denmark). Neurons were transfected in separate dishes with 0.5∼1 µg cDNA per coverslip. HEK293 cells were transfected with indicated plasmids using lipofectamine 3000 or 2000 according to manufacture instruction.

### Immunofluorescence staining and analysis

Neurons were fixed in 4 % paraformaldehyde/PBS for 20 min. Blocking and permeabilization was done in 4% dried milk/0.1 % Triton X-100/TBS. Primary and secondary antibodies were diluted in the same buffer and applied in separate incubation steps of 1 h and 45 min, respectively. Coverslips were mounted on glass microscope slides using ProLong Diamond anti-fade reagent (Thermo Fischer Scientific). We used the following primary antibodies (and dilutions): rat anti-total Tau (RTM38, serum at 1:5,000) raised against purified recombinant mouse Tau (Kubo et al., 2019), anti-GFP antibody (mouse monoclonal, 1:1,000, MBL, D153), anti-DHC1 (12345-1-AP, proteinteck), anti-AnkyrinG (1:1,000 mouse monoclonal, NeuroMAb), anti-acetylated tubulin (1:5,000, Sigma, 6-11B-1), anti-MAP2 antibody (1:1,000, Sigma, AP20), anti-pan-MAP2 (MAP2N, 1:1000) (Xie et al., 2014) and chicken anti-MAP2 (serum at 1:2,000, Biosensis).

### Live-cell imaging

For fluorescence recovery after photobleaching (FRAP) and photoconversion experiments, neurons expressing either GFP-tagged or Dendra2-tagged proteins were imaged using an Olympus FV-1000 microscope, Carl Zeiss LSM 700, or Leica SP8. Images were taken every 1 s. Photobleaching was induced by applying the 405 nm laser to a circular (3 µm diameter) spot at 100% laser power. Laser power was reduced to 2∼5% for photoconversion to avoid bleaching. Fluorescence intensity was measured using ImageJ, corrected for background fluorescence and the overall bleaching due to imaging, and normalized for the maximum and minimum fluorescence intensities.

### Dynein inhibitor

Cytoplasmic Dynein inhibitor, Ciliobrevin D, was purchased from calbiochem. Neurons were treated with 50 mM ciliobrevin D for 48 hours. Concentration of Ciliobrevin D, 50 mM, was based on previous literature that reported higher concentration, 100 mM, also inhibited Kinesin-1 and Kinesin-5 motor activity (Firestone et al., 2012). Effect of ciliobrevin D to localization of MAP2 was quantified by taking a ratio, signal intensity of distal axon (region after AIS) to proximal axon (region before AIS).

### Analysis of localization of MAP2C

#### MAP2c localization was analyzed as following

Signal intensities of one of dendrites and axon were measured in both green (endogenous) and red (exogenous) channels using image J. Selected area of axon is 168 ± 10 μm away from the cell body. Axon/dendrite ratio was calculated as following to normalize to endogenous MAP2 intensity in axon.

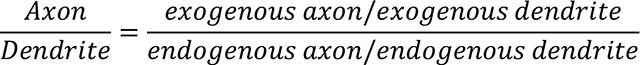

#### Immunoprecipitation

Cell lysates were prepared from transfected HEK293 cells 2 days after transfection. Cells were lysed with lysis buffer, 50 mM Hepes (pH 7.4), 100 mM NaCl, 2 mM EDTA, 0.5% Triton X-100, 1 mM NaF and protease inhibitor cocktail (Roche, 04693132001). After centrifuge at 4 ℃, 15,000 x g for 10 minutes, supernatants were collected as cell lysates. Protein concentration was determined by protein assay BCA kit (Nacalai, 06385-00). Immunoprecipitation was performed using Dynabeads protein G (invitrogen) according to provided manual. Briefly, anti-His antibody (D291-3, MBL) was incubated with Dynabeads protein G to bind antibody and add 500 μg lysate to antibody bound beads. After 1 hour incubation at 4 ℃, beads were washed with lysis buffer without NaF and protease inhibitor cocktail 3 times. Beads were transferred to new tubes and eluted with 50 μl of 1 x sample buffer. Immunoprecipitated DHC1 was quantified using Image J.

Mouse brain lysate was prepared and subjected to co-immunoprecipitation experiment as described below. Three-month-old female mouse was sacrificed by cervical dislocation and brain tissue was dissected. Dissected brain was homogenized with a glass-teflon homogenizer in ice-cold microtubule stabilizing buffer, MSB; 0.1 M MES (pH6.8), 10% glycerol, 1mM MgSO4, 1mM EGTA, 0.1 mM DTT, 0.5% triton X-100, 10 mM taxol, 2mM GTP, 0.1 mM PMSF, 0.1 mM DIFP, 1 mg/ml pepstatin, 1 mg/ml antipain, 10 mg/ml aprotinin, 10 mg/ml leupeptin, 50 µg/ml TLCK, x 1 protease inhibitor cocktail, 1 mM NaF, 1 mM β-glycerophosphatase, 1 mM Na3VO4, 0.5 µM okadaic acid. Brain lysate was prepared by removing cell debris by centrifuge at 2,400 x g for 3 minutes at 4℃ and subjected to coimmunoprecipitation. To perform co-immunoprecipitation experiment, anti-MAP2 (MAP2N) and control rabbit IgG was incubated with dynabeads protein G for 20 minutes at room temperature. Antibody-bound beads were incubated with 400 ml brain lysate and 100 ml MSB(-), MSB without phosphatase inhibitors and protease inhibitors, for 30 minutes at 4℃. After washing beads with MSB(-), co-IP samples were prepared by adding 1xSDS-PAGE samples buffer (2% SDS, 5% 2-mercaptoethanol, 10% glycerol, 62.5 mM Tris-HCl, pH6.8) to the beads.

Immunoreactive bands were detected with enhanced chemiluminescence (ECL, Perkin Elmer) and visualized by a detector (LAS3000, Fuji Film). Immunoreactivity was quantified using ImageJ.

#### Western blot

Proteins were separated in SDS polyacrylamide gels and electrophoretically transferred to nitrocellulose membranes. The membranes were washed, blocked with 4% non-fat dry milk in 150 mM NaCl, 10 mM sodium azide, and 20 mM Tris-HCl (pH8.0) for 30 min, and then incubated with primary antibodies overnight in the blocking buffer at 4°C. Membranes were incubated with following primary antibodies; anti-DHC1 (1:2000, Proteinteck, 12345-1-AP), anti-GFP (1:5000, MBL, 598), anti-MAP2 (1:1000, MAP2N). The membranes were washed and probed with HRP-conjugated secondary antibodies (KPL) according to the manufacturer’s instruction. Bands were detected with Immobilon Crescendo Western HRP substrate (Millipore, WBLUR0500) using LAS- 4000.

#### Statistical analyses

All statistical analyses were performed on GraphPad Prism software (GraphPad Software, Inc. CA, USA). Power analysis was performed using G*Power (Faul et al., 2007; Faul et al., 2009) with parameters taken from previous reports or similar experiments.

#### Simulation

All simulations were performed on the Virtual Cell (VCell) platform. Models will be publicly available in the VCell database upon publication.

We started building models with estimating the concentrations of MTs (75 µM), endogenous MAP2 (5 µM), and expressed GFP-MAP2 (25 µM) based on literatures and immunofluorescence data as described in Results. Then, a cylindrical compartment (2 µm diameter, 200 µm long), in which GFP-MAP2 freely diffuses, was constructed. GFP-MAP2 in a 3 µm spot was rapidly “bleached” by converting it to non-fluorescent MAP2 at a middle of the cylinder to mimic FRAP experiments. No flux was allowed at the boundary. The time evolution of GFP-MAP2 concentration (c) was described as the diffusion equation with a constant diffusion coefficient (*D*).

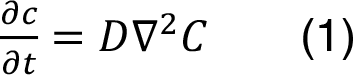

Simulation was performed using the fully implicit finite volume solver with varying time steps (max 100 ms). The concentration of GFP-MAP2 in the spot was measured and plotted. To determine the diffusion coefficient of GFP-MAP2, we took advantage of our extensive FRAP data on Tau lacking the MT-binding domain, which we found virtually freely diffusible, and free GFP. The virtual recovery curves were fitted with the experimental data. We changed the diffusion coefficient and monitored how the sum of squares changed. This allowed us to obtain diffusion coefficient providing the local minima in the sum of squares. The diffusion co-efficient of the GFP-MAP2 was adjusted to match the data of GFP-Tau and set to 10 µm^2^/s.

Next, we implemented MT-binding reactions of endogenous MAP2 and GFP-MAP2. These reactions are reversible and generate bound species for each. MT and bound species were virtually non-diffusible with a diffusion coefficient of 0.001 µm^2^/s. The initial condition, such as the concentration of MT-bound GFP-MAP2, was computed using a non-spatial simulation in a homogeneous environment. Then, to determine the binding kinetics, the virtual FRAP experiment was performed as described above. Here, the time evolution of GFP-MAP2 concentration was described as the reaction-diffusion equation.

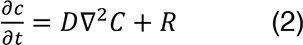

For the MT binding reaction of MAP2,

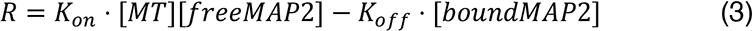

For this reaction, we set the initial Kon at 1 µM^-1^s^-1^, and Koff was adjusted to fit to the FRAP data of GFP-MAP2 to minimize the sum of squares. Then, Kon was changed from 10 to 0.01 µM^-1^s^-1^, and best fit Kon and Koff were searched. Kon larger than 1 µM^-1^s^-1^ provided Koff with similar fit to the values obtained at 1 µM^-1^s^-1^. The minimum Kon, which provided a fit as good as with 1 µM^-1^s^-1^ or higher, was around 0.1 µM^-1^s^-1^.

To simulate the cellular transport and localization of MAP2, we replaced the cylindrical geometry to a simple model neuron. Also, instead of photobleaching, photoconversion was implemented as a conversion of green MAP2 to red MAP2 species. Furthermore, a motor species was introduced to the model to realize active transport. MAP2 species could react with MT, motor, or MT-bound motor. Only MAP2-motor-MT was allowed to undergo directional transport at 1 µm/s. The time evolution of concentrations was described as the reaction-diffusion equation, except for the MAP2-motor-MT species, of which changes were described as the reaction-diffusion-advection equation with a constant velocity of transport (V).

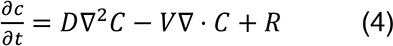

The initial condition was computed using a non-spatial simulation in a homogeneous environment. The initial concentrations of MT, motor, and motor-MT was set as constants to avoid the migration of these species over time by the transport (advection). MAP2 species were situated only in the soma and dendrites at the beginning. For the degradation model, irreversible reactions were implemented for MAP2 production and degradation with seed and remnant species.

For the stochastic simulation, we used Smoldyn, a particle-based simulator, in VCell with the parameters described above. A small cylindrical portion (5 µm x 0.2 µm x 0.2 µm) of a dendrite was simulated with a 0.01 s time step for 5 s using initial value boundary conditions. Dwell times of MAP2 on MT and motor-MT were analyzed using virtual kymographs for each species.

## Acknowledgement

We thank Drs. Stephanie Kaech and Yoko Toyoshima for kindly providing cDNA plasmids. We also thank Mr. Hirotaka Kishiue for technical assistance. This work was supported in part by a Grant- in-Aid for Scientific Research(C) JSPS KAKENHI (22H02946 for TM), a Grant-in-Aid for Scientific Research on Innovative Areas titled “Brain Protein Aging and Dementia Control” from MEXT (26117004 to TM), Scholarship Fund for Young/Women Researchers from The promotion and Mutual Aid Corporation for Private Schools of Japan (to YY), JST SPRING (JPMJSP2129 to RN and YS), and JSPS Core-to-Core Program A Advanced Research Networks (to HM). The Virtual Cell is supported by NIH Grant R24 GM137787 from the National Institute for General Medical Sciences.

